# Different correlation patterns between EEG and memory after E/I balance adjustment in normal middle-aged mice and AD model mice

**DOI:** 10.1101/2020.12.07.414433

**Authors:** Zheng Zhou, Liane Li, Mengsi Duan, Huaying Sun, Hu Yi, Yu Fu

## Abstract

Alzheimer’s disease (AD) is correlated with brain atrophy, neuronal loss, neurotransmitter imbalance, and cognitive decline, which also occur in normal aging. Thus, comparing the differences between normal aging and AD is of interest in order to test the accelerated aging hypothesis of AD. An imbalance between excitatory/inhibitory (E/I) neurotransmission, especially in gamma-aminobutyric acid (GABA) inhibition dysfunction, is involved in both AD and normal aging. In the present study, we performed correlation analyses between electroencephalograms (EEGs) and memory in middle-aged (∼12 months old) wild-type mice (WT) and AD model mice (APP/PS1) after E/I balance adjustment via GABA_A_ agonist muscimol and antagonist bicuculline administration (0.1 mg/kg intraperitoneally). Specifically, EEGs of the hippocampus and prefrontal cortex were recorded during Y-maze performance. Overall, WT and AD mice showed different correlation patterns between EEG activity and behavioral memory performance. Significant correlations were observed in EEG activity across a wider range of frequency bands (2–100 Hz, except 4–8 Hz) in WT mice, but were mainly observed in low frequency bands (delta-theta, 2–8 Hz) in AD mice. In addition, muscimol and bicuculline treatment contributed to better brain function in AD mice; in contrast, bicuculline administration resulted in poorer brain function in WT mice. Thus, our study suggests that AD shows a distinct pattern of disrupted brain function, rather than accelerated aging. Importantly, this work reveals new insights into future AD treatment by influencing low-frequency EEG activity through E/I balance adjustment, thereby aiding cognitive recovery.

## Introduction

Alzheimer’s disease (AD) is an important societal health concern. As life expectancy increases, the morbidity of neurodegenerative diseases, such as AD, also rapidly increases. At present, the incidence of AD is 4% among individuals aged 65 and up to 40%–50% among individuals aged 85 (1, 2). Evidence suggests that brain senescence contributes to AD pathogenesis (3, 4), and a link exists between the aging process and disease development (5-7). As aging is an important risk factor for AD, it would be interesting to compare differences between AD and normal aging, and thus help clarify the neural mechanisms of AD.

Normal aging and AD are both associated with brain atrophy, neuronal loss, neurotransmitter dysfunction, impaired functional connectivity, and cognitive decline, although these issues are more pronounced in AD than in normal aging (8). For example, AD patients exhibit worse memory dysfunction and deficits in the early stages compared with normal aging (8). Therefore, whether AD represents an acceleration of the aging process has been a topic of great interest (8-10).

Accumulating evidence suggests that an imbalance in excitation and inhibition (E/I), including that of the glutamate and gamma-aminobutyric acid (GABA) systems, is involved in both AD and aging (8, 11). Indeed, normal aging is accompanied by degeneration of GABAergic inhibition (12), with recent evidence also suggesting that GABA system dysfunction is involved in AD neuropathology (13, 14). Furthermore, changes in GABAergic neurotransmitters are thought to be related to cognitive deficits in AD (15), and GABAergic drugs have been used for cognitive recovery in AD treatment (16).

We previously highlighted the potential use of low-dose GABA_A_ drugs to improve cognition and restore neural network activity in AD (17). Notably, GABA_A_ agonists and antagonists may adjust E/I balance differently, resulting in brain function measures distributed scatteredly. In the current study, based on these scattered distributions, we analyzed the relationship between behavioral cognitive performance and cortical electrical activity in normal and AD mice. Both the AD and normal mice used in this study were at an early stage of senescence, with the degree of senescence in the 12-month-old mice approximate to that of 42.5-year-old humans (18). Thus, our results should provide information on the similarities and differences between normal aging and AD.

## Methods

In the present study, electrophysiological recordings were used following previous experiments (17), in which EEGs were recorded from mice with GABA_A_ intervention during Y-maze task performance (***Fig. 1A***). The work revealed changes in EEG frequency activity and behavioral memory after GABA_A_ agonist and antagonist administration, as well as the role of GABA_A_ intervention in AD model mice (***Fig. 1B***). In the current study, we pooled all drug-group data together for correlation analysis between EEG and behavioral changes and focused on differences in the correlation patterns between normal middle-aged and AD mice (***Fig. 1C***). Please see Fu et al. (2019) (17) for detailed information regarding electrophysiological recordings in mice.

**Fig. 1.**
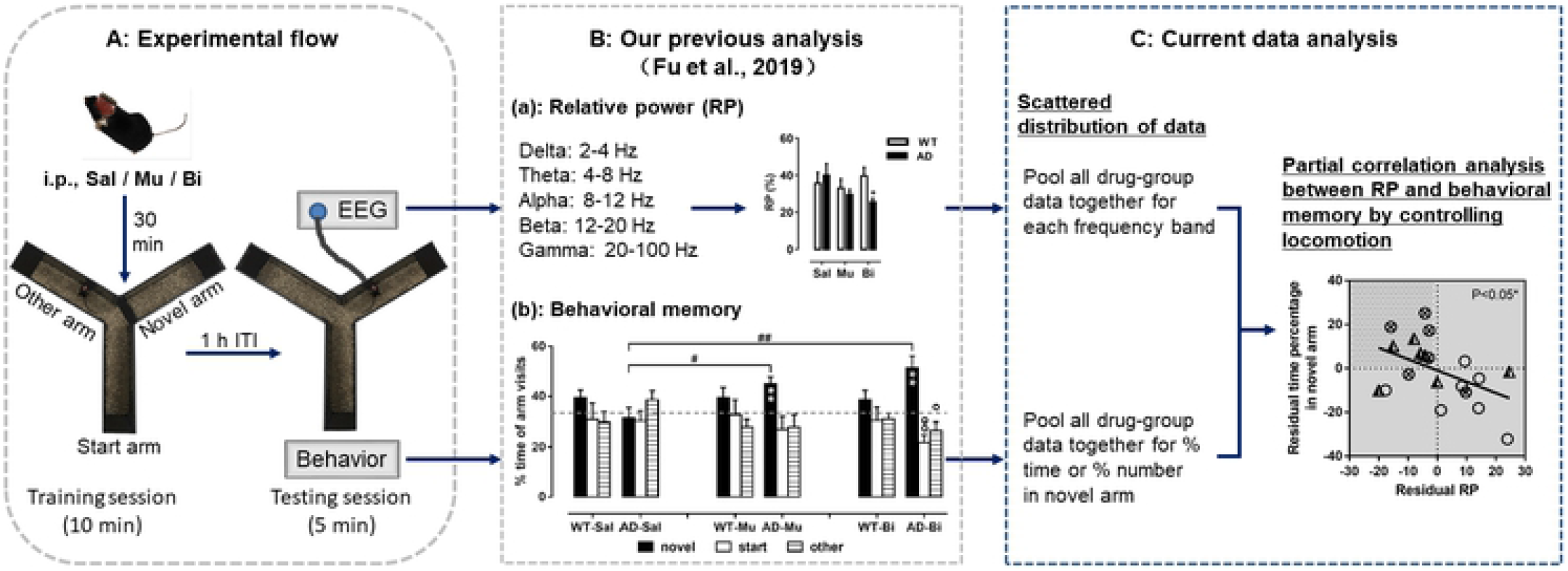
Experimental design and data analyses. (A) Experimental flow: animals were injected with GABA_A_ agonist (Mu), antagonist (Bi), or saline (Sal) 30 min before two-session Y-maze task, during which time EEGs were recorded. (B) Previous work showing EEG activity in five frequency bands (a, e.g.) and spatial recognition memory (b, e.g.), focusing on effects of GABA_A_ intervention in AD mice. (C) Current study showing correlation analysis of combined datasets between EEG activity (relative power; RP) and memory performance, focusing on differences in correlation patterns between normal middle-aged (WT) and AD mice.

### Animals and drugs

All animal and experimental care procedures complied with the guidelines for the National Care and Use of Animals and were approved by the National Animal Research Authority. APP/PS1 double transgenic male mice (amyloid precursor protein/presenilin-1 double transgenic, APPswe/PSEN1dE9; abbreviated to AD mice) and wild-type (WT) littermates were purchased from the Nanjing Biomedical Research Institute of Nanjing University (China). This mouse strain develops beta-amyloid (Aβ) deposits in the brain by 6–7 months of age. All mice included in this work were ∼12 months of age (53–55 weeks). They were housed in a temperature- and humidity-controlled room on a natural light-dark cycle with food and water available *ad libitum*. The animals were anesthetized with pentobarbital and intracardially perfused with saline (60–100ml) followed by 4% paraformaldehyde (100–200ml; Tianjin Guangfu Fine ChemicalResearch Institute, China). The brains were taken out and frozen sectioned at a 30-μm thickness to confirm the electrode locations.

The GABA_A_ receptor agonist muscimol hydrobromide (Sigma-Aldrich, St. Louis, MO, USA) and antagonist (+)-bicuculline (Selleck, Houston, TX, USA) were dissolved in 0.9% saline (Shandong Kangning Pharmaceutical Co., Ltd., China). In total, 22 AD mice and 19 WT mice were separately divided into three drug groups with 6–8 mice in each group. Mice were intraperitoneally (i.p.) injected with a single dose (0.1 mg/kg, 0.1 ml/10 g body weight) of muscimol (Mu) or bicuculline (Bi). Control mice received saline only. Drug administration was performed 30 min before the Y-maze task (***Fig. 1A***).

### EEG experiment

Before the EEG experiment, mice were implanted with three recording electrodes over the right hippocampus (Hip: AP: - 1.82 mm, ML: + 1.5 mm, DV: - 1.7 mm from skull), prefrontal cortex (PFC: AP: + 2.95 mm, ML: + 1.5 mm, DV: -0.75 mm from dura), and left occipital cortex (Ctx, as a control: AP: - 2.80 mm, ML: - 2.25 mm), respectively. The implantation surgery was conducted under pentobarbital anesthesia (60 mg/kg, i.p.; dissolved in 0.9% sodium, 10 mg/ml, Merck, Darmstadt, Germany). The Hip and PFC recording electrodes consisted of two twisted pairs of perfluoroalkoxy (PFA)-coated stainless steel wires (diameter 0.002’’, A-M systems, WA, USA), and that in the Ctx was a stainless-steel watch screw (M1.0 × L2.0 mm, RWD). In addition, two watch screws were placed in contact with the dura, one above the left olfactory bulb as a reference electrode and one in the central cerebellum as a ground electrode. All five electrodes were attached to a five-pin array and secured with dental acrylic. Animals were allowed at least one week to recover from surgery.

The Y-maze EEG recordings were performed in a shielding cage with a ceiling-mounted CCD camera to monitor behavior. The EEGs were recorded by a signal acquisition system (Intan Technologies, Los Angeles, CA, USA) consisting of an RHD2132 amplifier, RHD2000 USB interface board, and RHD2000 Interface GUI Software. The EEG sampling rate was 1 000 Hz. The five-pin array on the head of each mouse was connected to the amplifier, then to the interface board, and finally to the computer.

The Y-maze consisted of three arms (included angle of 120°) placed in the shielding cage. The walls of the cage and maze were decorated with a few visual spatial cues. The Y-maze task consisted of two sessions with a 1-h interval between them (***Fig. 1A***): (1) 10-min training session, in which the mouse was allowed to explore only two arms; (2) 5-min testing session, in which the mouse was allowed to explore all three arms. The arm blocked in the first session and open in the second session was novel to the animal, i.e., the novel arm.

### Data analysis

The EEG signals were analyzed off-line by MATLAB. Continuous EEG data were first separated into segments, each including 1 024 sample points. After rejection of segments with artifacts, each segment was filtered for five frequency bands: (1) delta: 2–4 Hz, (2) theta: 4–8 Hz, (3) alpha: 8–12 Hz, (4) beta: 12–20 Hz, and (5) gamma: 20–100 Hz. For each segment, the absolute power for each frequency band was calculated as: P = Σx^2^ / 1 024. Total power was the sum of the power of all frequency bands. The relative power (RP) for each frequency band was the percentage of its absolute power to total power. The mean RP of the testing session during the Y-maze task was used for further analysis.

Behavioral memory in each mouse was evaluated by the percentages of time spent in the novel arm (%Time) and number of entries in the novel arm (%Number) to all three arms during the Y-maze testing session. In addition, the total number of entries in all arms was used as an indicator of locomotion.

To investigate the correlation between EEG activity and behavioral memory, partial correlation analysis was used with locomotion as a covariate in the WT and AD groups. For each group, all drug data were pooled together for analysis. Partial correlation residuals were obtained by linear regression analysis, with EEG or behavioral memory as the dependent variables and locomotion as the independent variable. In the scatter plots of partial correlation residuals, the areas of interest (AOIs) were the upper-left corner for negative correlations (e.g., dotted area in ***Fig. 2A***) and upper-right corner for positive correlations (e.g., dotted area in ***Fig. 3A***). The animals distributed in these AOIs were those with better memory scores and decreased or increased EEG frequency activities. The ratios of the number of animals distributed in each AOI to total number of animals were compared among the different drug treatments by chi-squared tests. *P*-values of <0.05, <0.01, and< 0.001 were considered significant, highly significant, and very highly significant, respectively.

**Fig. 2.**
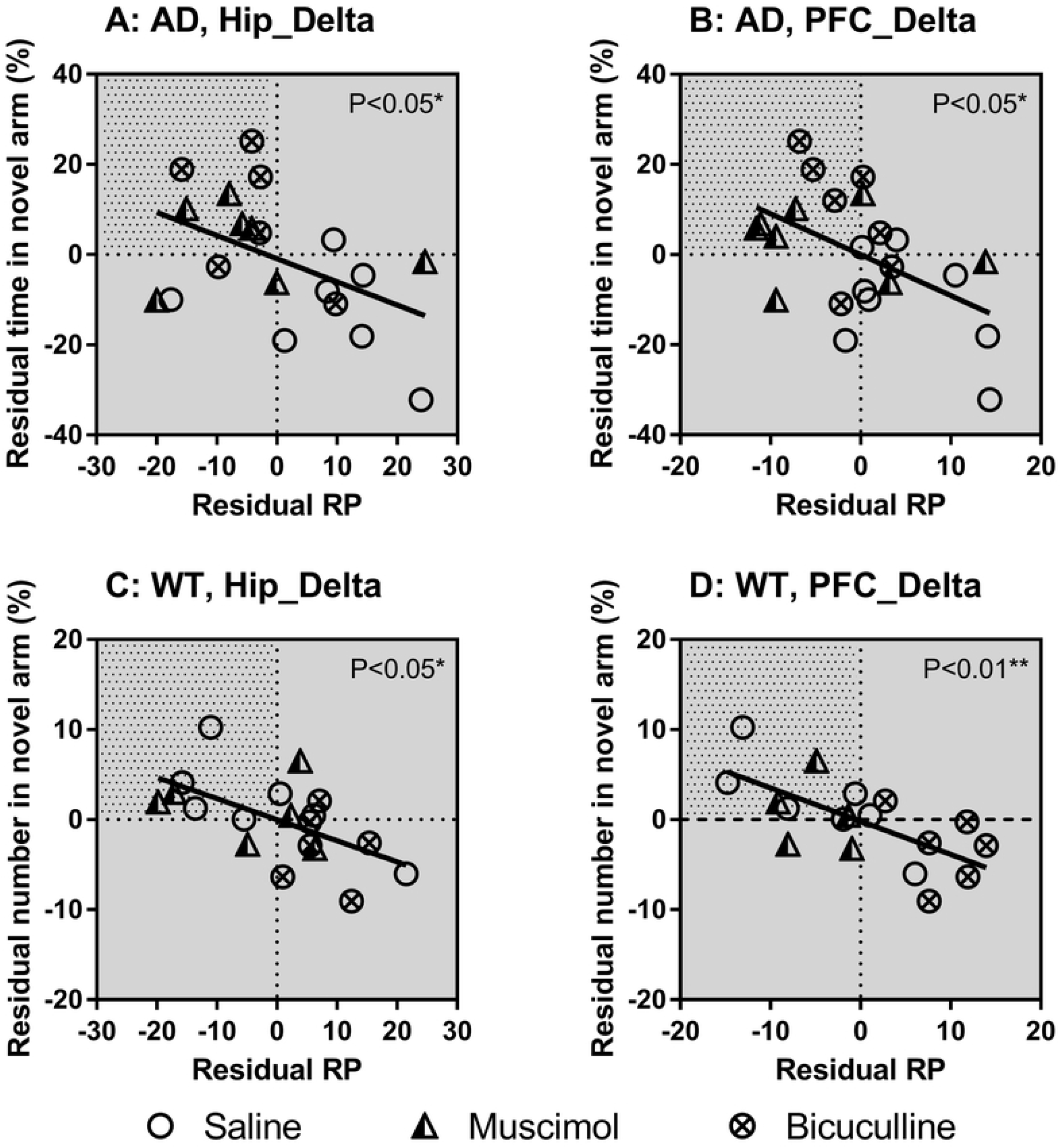
EEG activity in delta band showing negative correlations with memory performance in AD (A-B) and WT (C-D) mice. Animals distributed in area of interest (AOI, dotted area) showed better memory scores and decreased EEG activity. Partial correlation residuals were obtained by linear regression analysis. Abbreviations: RP, relative power; Hip, hippocampus; PFC, prefrontal cortex.

**Fig. 3.**
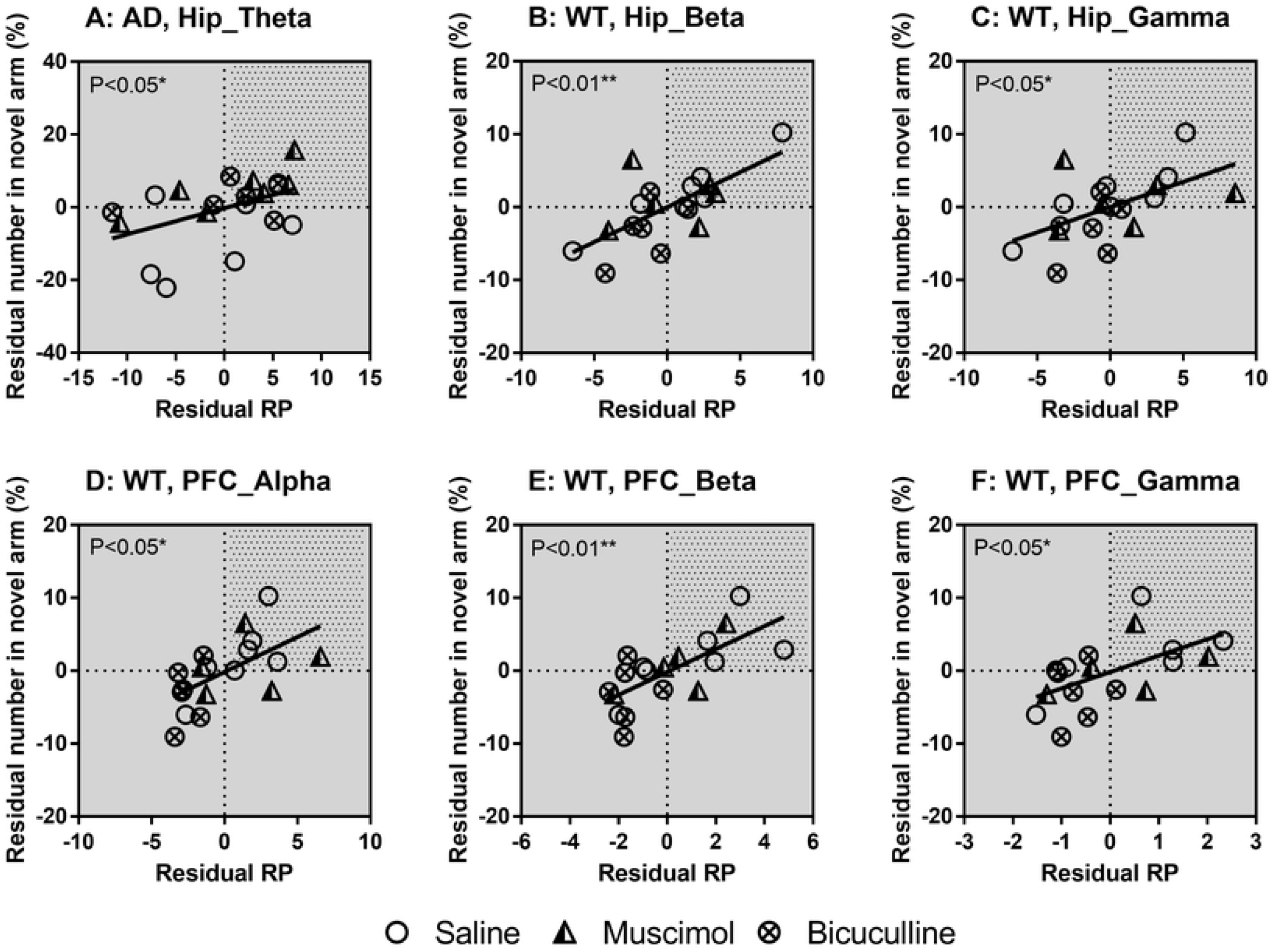
EEG activity in theta-gamma bands showing negative correlations with memory performance. (A) Hip EEG activity in theta band in AD mice; and (B-F) Hip/PFC EEG activity in alpha-gamma bands in WT mice. Animals distributed in AOI (dotted area) showed better memory scores and increased EEG activity. Partial correlation residuals were obtained by linear regression analysis. Abbreviations: RP, relative power; Hip, hippocampus; PFC, prefrontal cortex.

## Results

***Table 1*** displays the partial correlation analysis results between EEG activity and behavioral memory in the WT and AD mice. Results showed both similar and different correlations between the two groups. In both groups, Hip and PFC EEG activities in the delta band were significantly negatively correlated with time spent (%) or number of entries (%) in the novel arm (see ***Table 1*** for correlation coefficients and *P*-values, same below). In the WT group, Hip EEG activity in the beta-gamma bands and PFC EEG activity in the alpha-gamma bands were significantly positively correlated with number of entries (%) in the novel arm; in the AD group, Hip EEG activity in the theta band showed a significant positive correlation with the number of entries (%) in the novel arm. No significant correlations between Ctx EEG activity and behavioral memory were found in either group.

**Table 1.** Partial correlation analysis (controlling for locomotion) between EEG activity and memory performance in WT and AD mice.

| EEG<sup>a</sup> | Memory<sup>b</sup> | Hip |  |  |  | PFC |  |  |  | Ctx |  |  |  |
| --- | --- | --- | --- | --- | --- | --- | --- | --- | --- | --- | --- | --- | --- |
|  |  | WT |  | AD |  | WT |  | AD |  | WT |  | AD |  |
|  |  | *r* | *P* | *r* | *P* | *r* | *P* | *r* | *P* | *r* | *P* | *r* | *P* |
| *Delta* | % Time | -0.012 | 0.963 | **-0.474** | **0.040\*** | 0.112 | 0.668 | **-0.521** | **0.013\*** | 0.408 | 0.093 | 0.008 | 0.973 |
| *Theta* |  | -0.039 | 0.879 | 0.166 | 0.497 | -0.237 | 0.360 | 0.321 | 0.145 | -0.424 | 0.080 | 0.336 | 0.159 |
| *Alpha* |  | 0.172 | 0.495 | 0.300 | 0.211 | -0.129 | 0.622 | 0.358 | 0.102 | 0.242 | 0.333 | -0.015 | 0.952 |
| *Beta* |  | -0.049 | 0.847 | 0.376 | 0.113 | 0.259 | 0.316 | 0.322 | 0.144 | 0.325 | 0.189 | -0.358 | 0.133 |
| *Gamma* |  | -0.171 | 0.498 | 0.319 | 0.183 | 0.035 | 0.893 | 0.375 | 0.085 | 0.143 | 0.571 | -0.289 | 0.230 |
| *Delta* | % Number | **-0.580** | **0.012\*** | -0.353 | 0.138 | **-0.681** | **0.003\*\*** | -0.365 | 0.095 | -0.053 | 0.833 | 0.060 | 0.806 |
| *Theta* |  | -0.097 | 0.701 | **0.466** | **0.044\*** | 0.415 | 0.097 | 0.414 | 0.055 | -0.376 | 0.125 | 0.449 | 0.054 |
| *Alpha* |  | 0.427 | 0.077 | 0.110 | 0.654 | **0.587** | **0.013\*** | 0.293 | 0.186 | 0.240 | 0.337 | -0.282 | 0.243 |
| *Beta* |  | **0.693** | **0.001\*\*** | 0.142 | 0.561 | **0.701** | **0.002\*\*** | -0.025 | 0.911 | 0.431 | 0.074 | -0.162 | 0.509 |
| *Gamma* |  | **0.550** | **0.018\*** | 0.099 | 0.686 | **0.563** | **0.019\*** | 0.086 | 0.703 | 0.152 | 0.546 | -0.261 | 0.281 |

<sup>a</sup> EEG activity was calculated as relative power for each EEG frequency band ('Delta'-'Gamma').
<sup>b</sup> Memory performance was calculated as percentage of time spent ('% Time') or number of visits ('% Num') in novel arm of Y-maze.
Bold indicates significant differences in correlation: \* $P \leq 0.05$ and \*\* $P \leq 0.01$ .

The significant correlations above are also shown as scatter plots in ***Fig. 2-3***, with the partial correlation residuals for EEG and memory indices presented. The plots also show the distribution of animals injected with muscimol, bicuculline, or saline along the regression lines. Interestingly, different distributions were found between the WT and AD groups according to different drug administration, especially for bicuculline. For the negative correlations (***Fig. 2***), bicuculline treatment decreased Hip and PFC EEG activity in the delta band but improved memory performance in AD mice (***Fig. 2A-B***). In contrast, bicuculline treatment showed the opposite effect on WT mice (***Fig. 2C-D***). As seen in ***Table 2***, compared with the saline-treated mice, bicuculline administration significantly increased the number of AD animals in the AOI, but significantly decreased that number in the WT group (see ***Table 2*** for chi-square and *P*-values, same below). For the positive correlations (***Fig. 3***), bicuculline treatment mainly affected the WT mice, rather than the AD mice. In this group, drug treatment decreased Hip and PFC EEG activity in the mid-high frequency bands (alpha-gamma) and decreased memory performance (***Fig. 3B-F***). Furthermore, the number of animals in the AOI significantly decreased after bicuculline treatment compared with that after saline treatment (***Table 3, rows 2-6***).

**Table 2.** Comparison or number or analysis in area of in interest (AOI) among drug treatments for negetive correlations.

| | *Saline* | *Muscimol* | *Bicuculline* | $\chi^2_{Mu}$ | $P_{Mu}$ | $\chi^2_{Bi}$ | $P_{Bi}$ |
| --- | --- | --- | --- | --- | --- | --- | --- |
| *AD: Hip_Delta* | 0% (0/7) | 57% (4/7) | 67% (4/6) | 5.600 | ↑ **0.018\*** | 6.741 | ↑ **0.009\*\*** |
| *AD: PFC_Delta* | 0% (0/8) | 50% (4/8) | 43% (3/7) | 5.333 | ↑ **0.021\*** | 4.286 | ↑ **0.038\*** |
| *WT: Hip_Delta* | 57% (4/7) | 33% (2/6) | 0% (0/6) | 0.737 | 0.391 | 4.952 | ↓ **0.026\*** |
| *WT: PFC_Delta* | 71% (5/7) | 60% (3/5) | 0% (0/6) | 0.171 | 0.679 | 6.964 | ↓ **0.008\*\*** |

Please refer to upper left quadrants (dotted areas) of Fig. 2.
Bold indicates significant differences: \* $P \leq 0.05$ and \*\* $P \leq 0.01$ .

**Table 3.** Comparison of number of animals in area of interest (AOI) among drug treatments for positive correlations.

| | *Saline* | *Muscimol* | *Bicuculline* | $\chi^2_{Mu}$ | $P_{Mu}$ | $\chi^2_{Bi}$ | $P_{Bi}$ |
| --- | --- | --- | --- | --- | --- | --- | --- |
| *AD: Hip_Theta* | 29% (2/7) | 57% (4/7) | 50% (3/6) | 1.167 | 0.280 | 0.627 | 0.429 |
| *WT: Hip_Beta* | 71% (5/7) | 33% (2/6) | 0% (0/6) | 1.887 | 0.170 | 6.964 | ↓ **0.008\*\*** |
| *WT: Hip_Gamma* | 57% (4/7) | 33% (2/6) | 0% (0/6) | 0.737 | 0.391 | 4.952 | ↓ **0.026\*** |
| *WT: PFC_Alpha* | 71% (5/7) | 40% (2/5) | 0% (0/6) | 1.185 | 0.276 | 6.964 | ↓ **0.008\*\*** |
| *WT: PFC_Beta* | 57% (4/7) | 40% (2/5) | 0% (0/6) | 0.343 | 0.558 | 4.952 | ↓ **0.026\*** |
| *WT: PFC_Gamma* | 57% (4/7) | 40% (2/5) | 0% (0/6) | 0.343 | 0.558 | 4.952 | ↓ **0.026\*** |

Please refer to upper right quadrants (dotted areas) of Fig. 3.
Bold indicates significant differences: \* $P \leq 0.05$ and \*\* $P \leq 0.01$ .

In addition, muscimol treatment decreased Hip and PFC EEG activity in the delta band and improved memory in AD mice but not in WT mice (***Fig. 2-3***). In addition, the number of animals in the AOI was significantly higher after muscimol treatment than that after saline treatment (***Table 2, rows 1-2***).

## Discussion

In this study, we found that: (1) WT and AD mice showed different correlation patterns between Hip/PFC EEG activity and behavioral memory; (2) based on regression correlations, treatment with the GABA_A_ antagonist bicuculline improved brain function in AD mice, but resulted in poorer brain function in WT mice; (3) treatment with the GABA_A_ agonist muscimol benefited AD mice, but showed no obvious effects on WT mice.

Previous research has suggested that EEG oscillations in specific frequency bands can reflect behavioral or cognitive states (19). In the present study, correlations between Hip/PFC EEG activity in various frequency bands and behavioral memory differed in WT and AD mice. Correlations in EEG activity were detected across many frequency bands (except theta) in WT mice but were mainly found in the low-frequency bands (delta-theta) in AD mice. Because the WT and AD mice were performing the same task, differences in the behavior-EEG correlations may indicate differences in Hip/PFC functional state between them. Our results are similar to those of other studies. For example, earlier research showed that Hip-PFC connection strength and information transfer efficiency increased in control animals during working memory tasks, but showed no change in Aβ-injected animals (20).

In the current study, we also found that cognitive performance in AD mice was associated with EEG activity in the delta-theta frequency bands, which is worthy of attention. EEG slowing, especially increases in delta power, is a typical phenomenon accompanying cognitive symptoms in the AD brain (19). In addition, disrupted theta oscillations are observed in rats following exposure to Aβ and in transgenic AD mice with Aβ pathology (19). Importantly, EEG changes in these two frequency bands may occur during the early stages of AD (21). Thus, we found that spatial memory deficits were significantly related to delta-theta EEG activity, which may provide a direction for future AD treatment and early intervention of other such diseases, e.g., decreasing delta and increasing theta EEG activity to aid cognitive recovery. Indeed, several current noninvasive interventions involve the gradual improvement of the functional state of the AD brain via transcranial alternating current stimulation (22) or visual/auditory stimulation (23) at a specific frequency band.

Interestingly, in our study, treatment with the GABA_A_ antagonist bicuculline had the opposite effect on the behavior-EEG correlations in WT and AD mice. Delta EEG activity was negatively correlated with memory performance in both groups. However, bicuculline treatment in AD mice decreased delta activity and improved memory performance, contributing to better brain function, whereas bicuculline treatment in WT mice showed the opposite pattern, leading to poorer brain function. The negative effects of bicuculline in WT mice were also supported by decreased alpha-gamma EEG activity and impaired memory performance.

On the one hand, we previously discussed that modulation of E/I balance can contribute to recovery of brain function in AD mice (17). The benefits of the GABA_A_ antagonist may relate to the restoration of synaptic plasticity, compensation for the dampening of hyperexcitation, and cooperation with the cholinergic system (24-27). On the other hand, WT mice were at an early stage of senescence. Aging can impair the GABAergic inhibitory system, leading to E/I imbalance (5). From this point of view, GABA antagonists may potentiate the E/I imbalance toward excitation in aged animals. This imbalance could cause changes in neural network activity (5) and impair different forms of memory function, including Hip/PFC-dependent memory (28). This may explain why exposure to bicuculline in the normal-aged WT mice resulted in changes in the behavior-EEG correlations and poorer brain function.

Our study also showed that administration of the GABA_A_ agonist muscimol improved the behavior-EEG correlations in AD mice but showed no significant effects on WT mice. The apparent paradox of both muscimol and bicuculline having similar effects on AD mice has been discussed previously (17). A possible explanation is that muscimol exhibits more potent action at GABA_C_ than GABA_A_ receptors, but both receptors may share opposing actions on memory formation. The benefits of muscimol in our AD mice may also relate to neuroprotective effects, inhibition of Aβ-induced cascade, and normalization of GABA expression (17). Unexpectedly, we found that muscimol, as a GABA_A_ agonist, had no significant effect on the normal middle-aged (WT) mice. As GABA-mediated inhibition degrades during old age, muscimol should be effective on old animals (29). The reason for this may be due to the small sample size, relatively low dose of administration, and relatively young middle-aged animals. It would be interesting to determine whether muscimol is effective in older animals by adjusting these variables in future studies.

Whether AD represents an acceleration of the aging process has been discussed previously (30). Here, because the normal-aged mice showed different behavior-EEG correlation patterns to AD mice, and because the GABA_A_ antagonist produced the opposite effects on the two groups, we suggest that AD exhibits distinct and unique patterns of disrupted brain functional state, rather than accelerated healthy aging. Our results are in accordance with several previous studies (8, 9, 30), which suggest that normal aged brains exhibit general changes, but AD brains manifest changes in specific areas (30).

In summary, we found different correlation patterns between EEG and memory after E/I balance adjustment in normal middle-aged mice and AD mice. This work provides new insights into future AD treatment based on low-frequency EEG activity to aid cognitive recovery.

## Acknowledgments

This work was supported by the National Natural Science Foundation of China [81760258, 81560221, 81703497, and 61972158], Yunnan Ten Thousand Talents Plan Young & Elite Talents Project, and Natural Science Foundation of Yunnan Province of China [2017FB125 and 2017FD067]. We thank Prof. Xiangyang Kong for help with the experiments. We also thank Prof. Yuanye Ma and Dr. Nanhui Chen (Kunming Institute of Zoology, Chinese Academy of Sciences) for supervision of this project and data analysis.

